# Anterior cingulate folding pattern is altered in autism spectrum disorder

**DOI:** 10.1101/2025.06.24.661411

**Authors:** Ethan H. Willbrand, Samira A. Maboudian, Jacob J. Ludwig, Kevin S. Weiner

**Author notes:** Correspondence: Kevin Weiner.

## Abstract

Neuroimaging research has identified focal differences in the cerebral cortex of individuals with autism spectrum disorder (ASD), particularly in the cortical folds (sulci) within higher-level association cortices. The present study investigated the sulcal patterning and morphology of the anterior cingulate cortex (ACC) in individuals with ASD compared to neurotypical (NT) individuals for the first time. We used neuroimaging data from 50 NT and 50 ASD participants. All participants were under 20 years old and male. The two groups were age-matched. Using established criteria and cortical reconstructions generated from each participant’s T1-weighted magnetic resonance imaging scans with FreeSurfer, we identified the defining sulcal feature of ACC, the variably present paracingulate sulcus (PCGS): its presence in the left and right hemispheres, and asymmetry in PCGS presence between hemispheres. Finally, multiple quantitative morphological features (length, depth, and cortical thickness mean and standard deviation) were extracted from the PCGS using FreeSurfer tools. Analyses revealed that NT participants were more likely to have asymmetrical PCGS patterns than ASD participants (controlling for age and scanner site). However, none of the quantitative morphological features differed between groups. These findings suggest the presence of a variation in the prenatal neurodevelopment of ACC in young males with ASD; however, further research is necessary to uncover the role of this observed difference in the pathogenesis of ASD. The present study also adds to the growing literature implicating variations in PCGS patterning as a trait marker across multiple disorders.

**Lay Summary:** This study found that young males with autism spectrum disorder (ASD) show less hemispheric asymmetry in the presence of a notoriously variable brain structure (paracingulate sulcus (PCGS)) compared to neurotypical individuals. Considering that this feature of the PCGS develops before birth, the reduced asymmetry may indicate focal differences in brain development in ASD. These findings further enhance our understanding of the neurodevelopmental characteristics of ASD and highlight growing findings indicating that the PCGS may be a useful transdiagnostic marker for various psychiatric conditions.

## Introduction

Autism spectrum disorder (ASD) is a neurodevelopmental condition characterized by social, communicative, and behavioral challenges manifesting early in life and persisting throughout adulthood (American Psychiatric Association, 2013). Ongoing work identifies lobular and regional differences in the cerebral cortex of ASD individuals compared to neurotypical (NT) individuals (Hazlett et al., 2006; Mei et al., 2023; Prigge et al., 2021; Yang et al., 2016). Building on these large-scale differences, recent research identifies fine-scale neuroanatomical variations in individual sulci between groups, which, in some cases, have behavioral relevance (Ammons et al., 2021; Auzias et al., 2014; Brun et al., 2016; Ramos Benitez et al., 2024; Watanabe et al., 2014). Given these findings, and that the majority (60-70%) of the human cerebral cortex is buried within sulci (Ramos Benitez et al., 2024; Van Essen, 2007; Vogt et al., 1995; Willbrand, Maboudian, et al., 2023; Zilles et al., 1988), investigating the morphology of sulci in additional cortical expanses not yet examined in ASD could provide further insights into the neuroanatomical underpinnings of ASD.

In the present study, we focused on the anterior cingulate cortex (ACC) for three key reasons. First, the ACC is a key hub for cognitive control, social cognition, and emotional regulation—domains frequently affected in ASD (Apps et al., 2016; Di Martino et al., 2009; Margulies et al., 2007). Second, anatomical and functional alterations in the ACC (which did not consider individual sulci) have been previously reported in ASD, including differences in cortical thickness, surface area, connectivity, and activity patterns, suggesting an involvement in ASD pathophysiology (Balsters et al., 2017; Chien et al., 2021; Kohli et al., 2021; Laidi et al., 2019; Zhou et al., 2016). Third, within ACC resides the paracingulate sulcus (PCGS), a neuroanatomical feature of particular interest as it presents with striking individual variability in presence and length across individuals that have been related to individual differences in cognition with translational applications (Garrison et al., 2015; Paus et al., 1996; Yucel et al., 2001).

Specifically, variability in the presence, absence, asymmetry, and morphology of the PCGS has been known since the late 1800s and has been widely studied since then (Amiez et al., 2018; Chi et al., 1977; Paus et al., 1996; Yucel et al., 2001 are a few widely cited papers). Importantly, variations in its presence have been shown to correlate with cognitive processes related to mentalizing and multiple executive functions (Cachia et al., 2021)—which are commonly disrupted in ASD (Bylemans et al., 2023; Hemmers et al., 2022). Variability in PCGS morphology has also been implicated in other neurodevelopmental and psychiatric disorders, including schizophrenia, bipolar, and obsessive-compulsive disorders (Cachia et al., 2021). Structural alterations in the PCGS have been associated with impaired working memory and hallucinations in schizophrenia (Clark et al., 2010; Fornito et al., 2006; Garrison et al., 2015). Finally, previous work identified reduced gray matter in the vicinity of the PCGS between ASD and NT individuals at the group-level (Abell et al., 1999); however this study did not consider individual-level differences in the presence of the PCGS.

Despite its multifaceted relevance, to our knowledge, the PCGS has yet to be studied at the individual level in ASD. Therefore, the present study sought to lay the groundwork for future studies examining the PCGS in ASD through exploring three primary aims: (i) to characterize PCGS incidence and morphology in ASD individuals, (ii), to assess whether PCGS incidence differs in ASD individuals compared to NT individuals, and (iii) to determine whether PCGS morphology differs between ASD and NT individuals. Secondarily, we sought to quantify and compare the amount of ACC buried in sulci between ASD and NT individuals, and in individuals with and without a PCGS.

## Methods

Full methodological details are available in the **Supplementary Materials**. Briefly, the neuroanatomical data for each participant was leveraged from Autism Brain Imaging Data Exchange (ABIDE; Di Martino et al., 2014). Given limitations of this dataset resulting from females being historically underdiagnosed for ASD and less present in the ABIDE sample, as well as the large developmental age range of the ABIDE samples (childhood through young adulthood), we specifically selected participants that were male and less than 20 years old. In total, we randomly selected 100 participants fitting these criteria (50 participants in each group).

We also ensured that the NT and ASD groups were age matched (NT: 10.02 ± 2.03 years old; ASD: 9.32 ± 2.73 years old; *t*(98) = -1.45, *p =* 0.15). T1-weighted MRI scans were obtained for each participant from which reconstructions of the cortical surfaces were generated using FreeSurfer (Dale et al., 1999; Fischl et al., 1999). The PCGS was defined on native space surfaces and classified based on standard criteria (**Figure 1**; **Supplementary Materials**; Amiez et al., 2018; Garrison et al., 2015; Harper et al., 2022; Ono et al., 1990; Willbrand et al., 2024).

**Figure 1.**
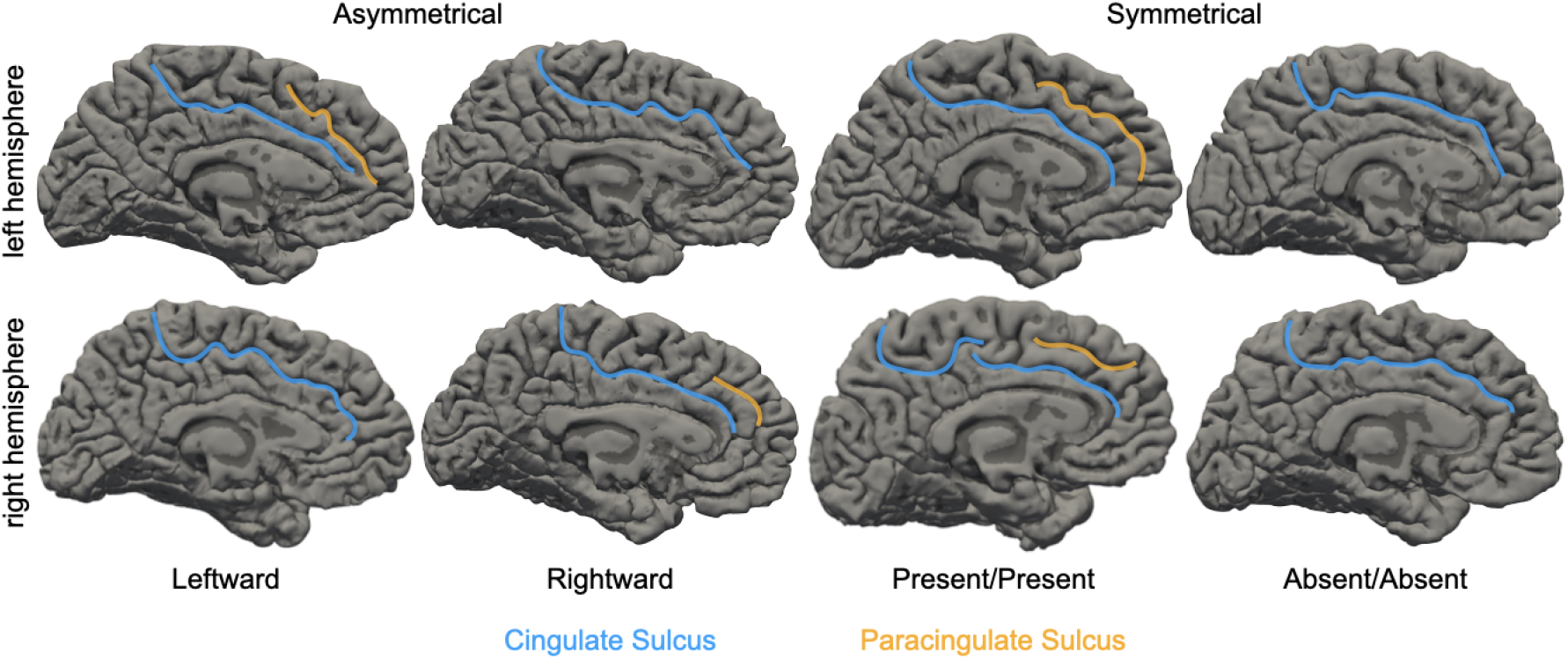
Examples of ACC Sulcal Patterning. ACC sulcal patterning is defined by the presence/absence of the PCGS, which runs dorsal and parallel to the CGS when present (CGS: blue; PCGS: orange). The images depict the intra-hemispheric presence/absence (rows) and inter-hemispheric asymmetry/symmetry (columns) of the PCGS in four example NT participants (columns) on the pial (i.e., wrinkled) FreeSurfer cortical surface reconstructions (Dale et al., 1999; Fischl et al., 1999). Sulci are dark gray and gyri are light gray. Abbreviations are as follows: anterior cingulate cortex (ACC), cingulate sulcus (CGS), paracingulate sulcus (PCGS).

All statistical tests were implemented in R (Team, 2020).

## Results

### Interhemispheric asymmetry in paracingulate sulcus presence is less common in autism spectrum disorder

Consistent with other studies in neurodevelopmental and clinical samples (Cachia et al., 2021), we observed prominent variability in PCGS presence within and between hemispheres (**Figure 2**). In both groups, the PCGS was more often present than absent in the left hemisphere (NT: χ*2*(1) = 13.52, *p* = 0.00023; ASD: χ*2*(1) = 5.12, *p* = 0.023), but not the right hemisphere (NT: χ*2*(1) = 0.32, *p* = 0.57; ASD: χ*2*(1) = 0.08, *p* = 0.77; **Figure 2A**). The presence of a PCGS was more frequent in the left than right hemisphere in NT participants (χ*2*(1) = 5.32, *p* = 0.021), but not in ASD participants (χ*2*(1) = 2.03, *p* = 0.15; **Figure 2A**). In addition, implementing McNemar’s test for symmetry revealed significant asymmetry for NT participants but not for ASD participants (NT: χ*2*(1) = 4.48, *p* = 0.034; ASD: χ*2*(1) = 3.27, *p* = 0.07; **Figure 2B**).

**Figure 2.**
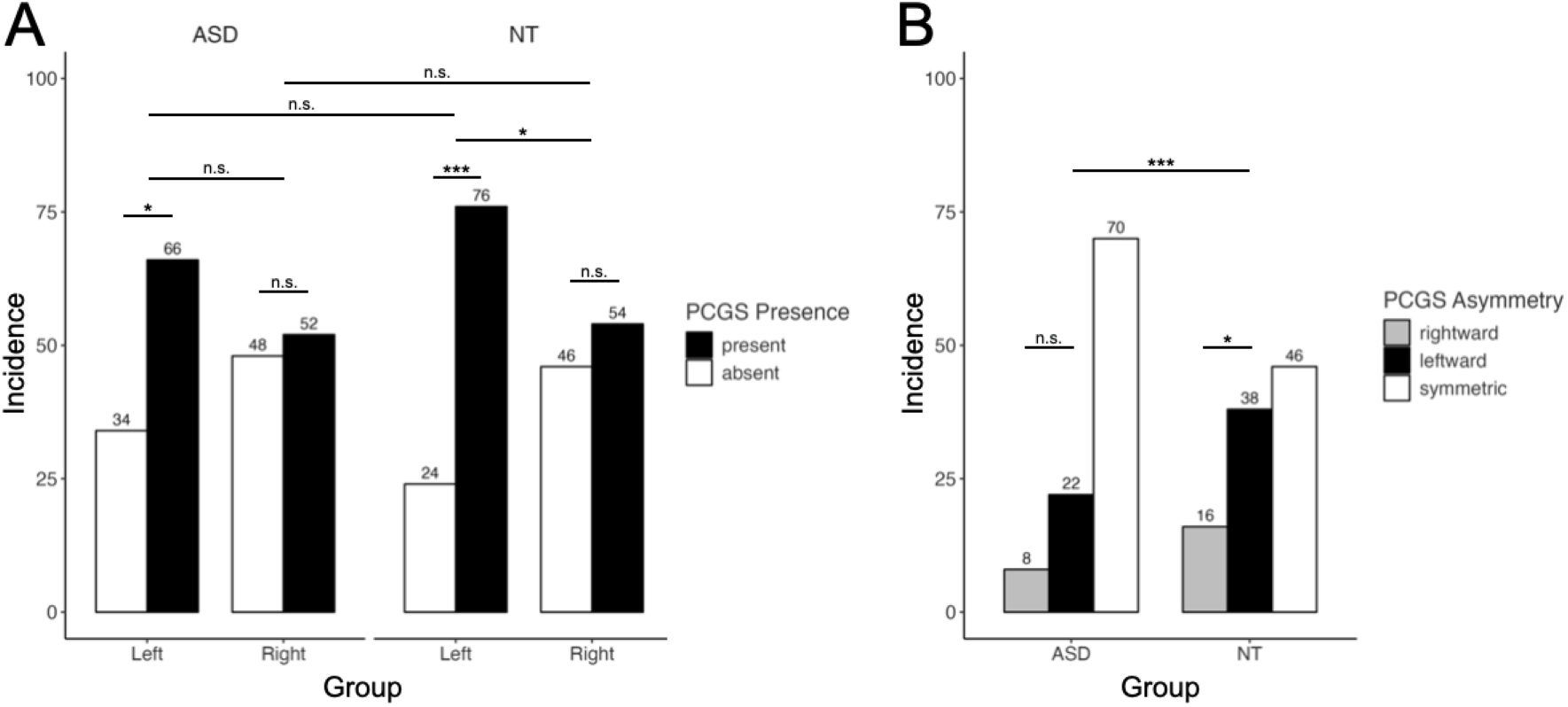
Differences in PCGS hemispheric incidence and asymmetry between NT and ASD individuals. **(A)** Bar plot visualizing PCGS incidence as a function of hemisphere (x-axis), group, and presence (colors; see key). **(B)** Bar plot visualizing PCGS incidence as a function of group (x-axis) and asymmetry (colors; see key). Upper asterisk denotes GLM results for between-group comparisons of asymmetry, while lower asterisks depict McNemar test results for within-group comparisons of asymmetry. In both (A) and (B), the percentages are included above each column (*n* = 50 in each group). Abbreviations are as follows: autism spectrum disorder (ASD), group, neurotypical (NT), paracingulate sulcus (PCGS). Asterisks indicate the following *p-*value thresholds: *** *p* < 0.001, * *p* < 0.05, n.s. *p* > .05.

Next, we explicitly compared PCGS presence and asymmetry between the two groups by implementing logistic regression GLMs with a predictor of group (controlling for age and scanner site). We observed that ASD participants were not less likely than NT participants to have a PCGS in either hemisphere (no main effect of group; Left: χ*2*(1) = 1.35, *p* = 0.24; Right: χ*2*(1) = 0.23, *p* = 0.62; **Figure 2A**). However, there was a main effect of group on PCGS asymmetry (χ*2*(1) = 10.83, *p* = 0.0009), which showed that NT participants had higher odds of having an asymmetric PCGS compared to ASD participants (OR [95% CI] = 4.55 [1.82-12.23]; bootstrapped OR [95% CI] = 3.02 [1.30-7.74]; **Figure 2B**). Quantitative morphological features of the PCGS (length, depth, and CT mean and STD) and the amount of ACC buried in sulci did not differ between groups (**Supplementary Materials**).

## Discussion

Here, we comprehensively examined PCGS features in young male NT and ASD individuals.

Our findings revealed significant differences in PCGS asymmetry between groups, with NTs exhibiting more pronounced asymmetry than those with ASD. This finding aligns with previous ASD research examining sulci in other parts of the brain (Ammons et al., 2021; Brun et al., 2016; Ramos Benitez et al., 2024; Watanabe et al., 2014). The prenatal emergence of sulci (e.g., the PCGS appears around 36 gestational weeks; Chi et al., 1977; Nishikuni & Ribas, 2013; Welker, 1990) suggests the differences observed in ASD in the present study may arise from early disruptions in genetic factors, cellular mechanisms, and biomechanical forces that underlie the formation of sulci (Borrell, 2018; Kroenke & Bayly, 2018). Further studies using fetal imaging or postmortem analyses could help elucidate mechanisms leading to altered sulcal patterns in ASD.

The observed asymmetry in NT, but not ASD, participants may have implications for understanding the neural basis of cognitive and behavioral differences associated with ASD. The PCGS has been linked to cognitive processes such as mentalizing and executive functions, which are often disrupted in ASD (Bylemans et al., 2023; Cachia et al., 2021; Hemmers et al., 2022). Further, the presence of the PCGS has been associated with variations in cytoarchitectonic (Amiez et al., 2021; Vogt et al., 1995), anatomical and functional connectivity (Fedeli et al., 2020; Harper et al., 2024), and task-based activation (Amiez et al., 2013; Amiez & Petrides, 2014; Artiges et al., 2006; Crosson et al., 1999). Therefore, the observed reduction in PCGS asymmetry in ASD may reflect alterations in the underlying anatomical and functional organization supporting these cognitive functions, which may contribute to the social and communicative challenges characteristic of ASD (Hemmers et al. 2022; Bylemans et al. 2023). This aligns with previous studies showing disrupted interhemispheric connectivity and atypical lateralization patterns in ASD—studies that crucially did not consider the PCGS (Cardinale et al., 2013; Floris et al., 2021). Further research using functional MRI or diffusion tensor imaging could help clarify the extent to which PCGS alterations impact broader network-level connectivity and behavioral outcomes in ASD.

Broadly, these findings add to growing research showing that PCGS morphology (left/right hemisphere absence/presence, asymmetry, length) may be markers for brain dysfunction in certain psychiatric (schizophrenia, bipolar, and obsessive-compulsive disorders; Cachia et al., 2021) and neurological (frontotemporal dementia; Harper et al., 2022, 2023) disorders. While many features of the PCGS are associated with multiple disorders, it is not a consistent one to all relationship (e.g., PCGS asymmetry is reduced in some disorders, but not all examined thus far; Cachia et al., 2021; Fornito et al., 2007; Le Provost et al., 2003; Meredith et al., 2012; Shim et al., 2009; Whittle et al., 2009; Yücel et al., 2002). Therefore, it is necessary for future investigations to continue testing the generalizability of which PCGS features are associated with which disorders, and, concomitantly, then determine what processes underlie their initial formation in utero and continued development after birth. Further, this observed regional specificity in sulcal changes suggests that different cortical areas may be differentially vulnerable to ASD-related neurodevelopmental disruptions. Future research should examine the interplay between these observed regional anatomical differences in ASD cortical morphology and different cognitive tasks.

In terms of limitations, future studies should consider a more diverse sample to fully capture the variability of the PCGS across the autism spectrum, examine the potential cognitive consequences of these variations, and include additional regions containing variable sulci (e.g., lateral prefrontal, ventromedial prefrontal, and lateral parietal cortices; Lopez-Persem et al., 2019; Willbrand et al., 2022; Willbrand, Tsai, et al., 2023) to examine more complex neuroanatomical interactions. Further development and validation of automated tools for tertiary sulcal identification (Borne et al., 2020, 2021; Hao et al., 2020; Lee et al., 2024, 2025)— especially for sulci not present in all hemispheres—would facilitate larger-scale investigations.

## Conclusion

This study is the first, to our knowledge, to examine PCGS features in ASD and identifies altered asymmetry in this tertiary sulcus as a potential neurodevelopmental marker. These findings emphasize the value of investigating sulcal morphology beyond traditional morphological metrics in ASD research and support further exploration of the PCGS in relation to behavior and development. As understanding of sulcal variability advances, so too will insights into the neurobiological substrates of ASD and other neurodevelopmental conditions.

## Acknowledgments

This research was supported by NSF CAREER Award 2042251 (Weiner), Medical Scientist Training Program Grant T32 GM140935 (Willbrand), and NIH F31 AG082446-01A1 (Maboudian). Funding for original data collection and curation for the New York University sample was provided by NIH (K23 MH087770; R21 MH084126; R01 MH081218; R01 HD065282), Autism Speaks, The Stavros Niarchos Foundation, The Leon Levy Foundation, and an endowment provided by Phyllis Green and Randolph Co□wen. Funding for original data collection and curation for the Georgetown University sample was provided by NIMH MH084961, Intellectual and Developmental Disabilities Research Center, and Children’s National Medical Center HD040677-07.

## Author contributions

EHW, SAM, and KSW designed research; EHW, SAM, JJL, and KSW performed research; EHW, SAM, JJL, and KSW wrote the paper; all authors gave final approval to the paper before submission.

## Data and code availability

All data and original code used for the present project will be publicly available on GitHub upon publication (https://github.com/cnl-berkeley/stable_projects). Any additional information required to reanalyze the data reported in this paper is available from the corresponding author (Kevin Weiner) upon request.

## References

Abell, F., Krams, M., Ashburner, J., Passingham, R., Friston, K. J., Frackowiak, R. S. J., Happé, F., Frith, C., & Frith, U. (1999). The neuroanatomy of autism: a voxel-based whole brain analysis of structural scans. Neuroreport, 10(8), 1647–1651. 10.1097/00001756-199906030-00005

American Psychiatric Association, D. (2013). academia.edu. https://www.academia.edu/download/38718268/csl6820_21.pdf

Amiez, C., Neveu, R., Warrot, D., Petrides, M., Knoblauch, K., & Procyk, E. (2013). The location of feedback-related activity in the midcingulate cortex is predicted by local morphology. The Journal of Neuroscience: The Official Journal of the Society for Neuroscience, 33(5), 2217–2228. 10.1523/JNEUROSCI.2779-12.2013

Amiez, C., & Petrides, M. (2014). Neuroimaging evidence of the anatomo-functional organization of the human cingulate motor areas. Cerebral Cortex, 24(3), 563–578. 10.1093/cercor/bhs329

Amiez, C., Sallet, J., Novek, J., Hadj-Bouziane, F., Giacometti, C., Andersson, J., Hopkins, W. D., & Petrides, M. (2021). Chimpanzee histology and functional brain imaging show that the paracingulate sulcus is not human-specific. Communications Biology, 4(1), 54. 10.1038/s42003-020-01571-3

Amiez, C., Wilson, C. R. E., & Procyk, E. (2018). Variations of cingulate sulcal organization and link with cognitive performance. Scientific Reports, 8(1), 1–13. 10.1038/s41598-018-32088-9

Ammons, C. J., Winslett, M.-E., Bice, J., Patel, P., May, K. E., & Kana, R. K. (2021). The mid-fusiform sulcus in autism spectrum disorder: Establishing a novel anatomical landmark related to face processing. Autism Research: Official Journal of the International Society for Autism Research, 14(1), 53–64. 10.1002/aur.2425

Apps, M. A. J., Rushworth, M. F. S., & Chang, S. W. C. (2016). The anterior cingulate gyrus and social cognition: Tracking the motivation of others. Neuron, 90(4), 692–707. 10.1016/j.neuron.2016.04.018

Artiges, E., Martelli, C., Naccache, L., Bartrés-Faz, D., LeProvost, J.-B., Viard, A., Paillère-Martinot, M.-L., Dehaene, S., & Martinot, J.-L. (2006). Paracingulate sulcus morphology and fMRI activation detection in schizophrenia patients. In Schizophrenia Research (Vol. 82, Issues 2-3, pp. 143–151). 10.1016/j.schres.2005.10.022

Auzias, G., Viellard, M., Takerkart, S., Villeneuve, N., Poinso, F., Fonséca, D. D., Girard, N., & Deruelle, C. (2014). Atypical sulcal anatomy in young children with autism spectrum disorder. NeuroImage. Clinical, 4, 593–603. 10.1016/j.nicl.2014.03.008

Balsters, J. H., Apps, M. A. J., Bolis, D., Lehner, R., Gallagher, L., & Wenderoth, N. (2017). Disrupted prediction errors index social deficits in autism spectrum disorder. Brain: A Journal of Neurology, 140(1), 235–246. 10.1093/brain/aww287

Borne, L., Rivière, D., Cachia, A., Roca, P., Mellerio, C., Oppenheim, C., & Mangin, J.-F. (2021). Automatic recognition of specific local cortical folding patterns. NeuroImage, 238, 118208. 10.1016/j.neuroimage.2021.118208

Borne, L., Rivière, D., Mancip, M., & Mangin, J.-F. (2020). Automatic labeling of cortical sulci using patch- or CNN-based segmentation techniques combined with bottom-up geometric constraints. Medical Image Analysis, 62, 101651. 10.1016/j.media.2020.101651

Borrell, V. (2018). How Cells Fold the Cerebral Cortex. The Journal of Neuroscience: The Official Journal of the Society for Neuroscience, 38(4), 776–783. 10.1523/JNEUROSCI.1106-17.2017

Brun, L., Auzias, G., Viellard, M., Villeneuve, N., Girard, N., Poinso, F., Da Fonseca, D., & Deruelle, C. (2016). Localized Misfolding Within Broca’s Area as a Distinctive Feature of Autistic Disorder. In Biological Psychiatry: Cognitive Neuroscience and Neuroimaging (Vol. 1, Issue 2, pp. 160–168). 10.1016/j.bpsc.2015.11.003

Bylemans, T., Heleven, E., Baetens, K., Deroost, N., Baeken, C., & Van Overwalle, F. (2023). Mentalizing and narrative coherence in autistic adults: Cerebellar sequencing and prediction. Neuroscience and Biobehavioral Reviews, 146(105045), 105045. 10.1016/j.neubiorev.2023.105045

Cachia, A., Borst, G., Jardri, R., Raznahan, A., Murray, G. K., Mangin, J.-F., & Plaze, M. (2021). Towards Deciphering the Fetal Foundation of Normal Cognition and Cognitive Symptoms From Sulcation of the Cortex. Frontiers in Neuroanatomy, 15, 712862. 10.3389/fnana.2021.712862

Cardinale, R. C., Shih, P., Fishman, I., Ford, L. M., & Müller, R.-A. (2013). Pervasive rightward asymmetry shifts of functional networks in autism spectrum disorder. JAMA Psychiatry (Chicago, Ill.), 70(9), 975–982. 10.1001/jamapsychiatry.2013.382

Chien, Y.-L., Chen, Y.-C., & Gau, S. S.-F. (2021). Altered cingulate structures and the associations with social awareness deficits and CNTNAP2 gene in autism spectrum disorder. NeuroImage. Clinical, 31(102729), 102729. 10.1016/j.nicl.2021.102729

Chi, J. G., Dooling, E. C., & Gilles, F. H. (1977). Gyral development of the human brain. Annals of Neurology, 1(1), 86–93. 10.1002/ana.410010109

Clark, G. M., Mackay, C. E., Davidson, M. E., Iversen, S. D., Collinson, S. L., James, A. C., Roberts, N., & Crow, T. J. (2010). Paracingulate sulcus asymmetry; sex difference, correlation with semantic fluency and change over time in adolescent onset psychosis. Psychiatry Research, 184(1), 10–15. 10.1016/j.pscychresns.2010.06.012

Crosson, B., Sadek, J. R., Bobholz, J. A., Gökçay, D., Mohr, C. M., Leonard, C. M., Maron, L., Auerbach, E. J., Browd, S. R., Freeman, A. J., & Briggs, R. W. (1999). Activity in the paracingulate and cingulate sulci during word generation: an fMRI study of functional anatomy. Cerebral Cortex, 9(4), 307–316. 10.1093/cercor/9.4.307

Dale, A. M., Fischl, B., & Sereno, M. I. (1999). Cortical surface-based analysis. I. Segmentation and surface reconstruction. NeuroImage, 9(2), 179–194. 10.1006/nimg.1998.0395

Di Martino, A., Ross, K., Uddin, L. Q., Sklar, A. B., Castellanos, F. X., & Milham, M. P. (2009). Functional brain correlates of social and nonsocial processes in autism spectrum disorders: an activation likelihood estimation meta-analysis. Biological Psychiatry, 65(1), 63–74. 10.1016/j.biopsych.2008.09.022

Di Martino, A., Yan, C.-G., Li, Q., Denio, E., Castellanos, F. X., Alaerts, K., Anderson, J. S., Assaf, M., Bookheimer, S. Y., Dapretto, M., Deen, B., Delmonte, S., Dinstein, I., Ertl-Wagner, B., Fair, D. A., Gallagher, L., Kennedy, D. P., Keown, C. L., Keysers, C., … Milham, M. P. (2014). The autism brain imaging data exchange: towards a large-scale evaluation of the intrinsic brain architecture in autism. Molecular Psychiatry, 19(6), 659–667. 10.1038/mp.2013.78

Fedeli, D., Del Maschio, N., Caprioglio, C., Sulpizio, S., & Abutalebi, J. (2020). Sulcal Pattern Variability and Dorsal Anterior Cingulate Cortex Functional Connectivity Across Adult Age. Brain Connectivity, 10(6), 267–278. 10.1089/brain.2020.0751

Fischl, B., Sereno, M. I., & Dale, A. M. (1999). Cortical surface-based analysis. II: Inflation, flattening, and a surface-based coordinate system. NeuroImage, 9(2), 195–207. 10.1006/nimg.1998.0396

Floris, D. L., Wolfers, T., Zabihi, M., Holz, N. E., Zwiers, M. P., Charman, T., Tillmann, J., Ecker, C., Dell’Acqua, F., Banaschewski, T., Moessnang, C., Baron-Cohen, S., Holt, R., Durston, S., Loth, E., Murphy, D. G. M., Marquand, A., Buitelaar, J. K., Beckmann, C. F., & EU-AIMS Longitudinal European Autism Project Group. (2021). Atypical brain asymmetry in autism-A candidate for clinically meaningful stratification. Biological Psychiatry: Cognitive Neuroscience and Neuroimaging, 6(8), 802–812. 10.1016/j.bpsc.2020.08.008

Fornito, A., Malhi, G. S., Lagopoulos, J., Ivanovski, B., Wood, S. J., Velakoulis, D., Saling, M. M., McGorry, P. D., Pantelis, C., & Yücel, M. (2007). In vivo evidence for early neurodevelopmental anomaly of the anterior cingulate cortex in bipolar disorder. Acta Psychiatrica Scandinavica, 116(6), 467–472. 10.1111/j.1600-0447.2007.01069.x

Fornito, A., Yücel, M., Wood, S. J., Proffitt, T., McGorry, P. D., Velakoulis, D., & Pantelis, C. (2006). Morphology of the paracingulate sulcus and executive cognition in schizophrenia. Schizophrenia Research, 88(1-3), 192–197. 10.1016/j.schres.2006.06.034

Garrison, J. R., Fernyhough, C., McCarthy-Jones, S., Haggard, M., Australian Schizophrenia Research Bank, & Simons, J. S. (2015). Paracingulate sulcus morphology is associated with hallucinations in the human brain. Nature Communications, 6, 8956. 10.1038/ncomms9956

Hao, L., Bao, S., Tang, Y., Gao, R., Parvathaneni, P., Miller, J. A., Voorhies, W., Yao, J., Bunge, S. A., Weiner, K. S., Landman, B. A., & Lyu, I. (2020). Automatic Labeling of Cortical Sulci Using Spherical Convolutional Neural Networks in a Developmental Cohort. 2020 IEEE 17th International Symposium on Biomedical Imaging (ISBI), 412–415. 10.1109/ISBI45749.2020.9098414

Harper, L., de Boer, S., Lindberg, O., Lätt, J., Cullen, N., Clark, L., Irwin, D., Massimo, L., Grossman, M., Hansson, O., Pijnenburg, Y., McMillan, C. T., & Santillo, A. F. (2023). Anterior cingulate sulcation is associated with onset and survival in frontotemporal dementia. Brain Communications, 5(5), fcad264. 10.1093/braincomms/fcad264

Harper, L., Lindberg, O., Bocchetta, M., Todd, E. G., Strandberg, O., van Westen, D., Stomrud, E., Landqvist Waldö, M., Wahlund, L.-O., Hansson, O., Rohrer, J. D., & Santillo, A. (2022). Prenatal Gyrification Pattern Affects Age at Onset in Frontotemporal Dementia. Cerebral Cortex. 10.1093/cercor/bhab457

Harper, L., Strandberg, O., Spotorno, N., Nilsson, M., Lindberg, O., Hansson, O., & Santillo, A. F. (2024). Structural and functional connectivity associations with anterior cingulate sulcal variability. Research Square. 10.21203/rs.3.rs-3831519/v1

Hazlett, H. C., Poe, M. D., Gerig, G., Smith, R. G., & Piven, J. (2006). Cortical gray and white brain tissue volume in adolescents and adults with autism. Biological Psychiatry, 59(1), 1–6. 10.1016/j.biopsych.2005.06.015

Hemmers, J., Baethge, C., Vogeley, K., & Falter-Wagner, C. M. (2022). Are executive dysfunctions relevant for the autism-specific cognitive profile? Frontiers in Psychiatry, 13, 886588. 10.3389/fpsyt.2022.886588

Kohli, J., Martindale, I., Wilkinson, M., Kinnear, M., Linke, A., Hau, J., Mueller, R.-A., & Carper, R. (2021). Altered links between anterior cingulate cortex anatomy and inhibition skills in middle to older aged adults with autism spectrum disorder (ASD). Biological Psychiatry, 89(9), S270–S271. 10.1016/j.biopsych.2021.02.677

Kroenke, C. D., & Bayly, P. V. (2018). How Forces Fold the Cerebral Cortex. The Journal of Neuroscience: The Official Journal of the Society for Neuroscience, 38(4), 767–775. 10.1523/JNEUROSCI.1105-17.2017

Laidi, C., Boisgontier, J., de Pierrefeu, A., Duchesnay, E., Hotier, S., d’Albis, M.-A., Delorme, R., Bolognani, F., Czech, C., Bouquet, C., Amestoy, A., Petit, J., Holiga, Š., Dukart, J., Gaman, A., Toledano, E., Ly-Le Moal, M., Scheid, I., Leboyer, M., & Houenou, J. (2019). Decreased cortical thickness in the anterior cingulate cortex in adults with autism. Journal of Autism and Developmental Disorders, 49(4), 1402–1409. 10.1007/s10803-018-3807-3

Lee, S., Lee, S., Willbrand, E. H., Parker, B. J., Bunge, S. A., Weiner, K. S., & Lyu, I. (2024). Leveraging input-level feature deformation with guided-attention for sulcal labeling. IEEE Transactions on Medical Imaging, PP(99), 1–1. 10.1109/TMI.2024.3468727

Lee, S., Son, J., Lee, S., Willbrand, E. H., Parker, B. J., Weiner, K. S., & Lyu, I. (2025). Extensive spherical region enlargement with isotropic deformation for sulcal labeling. 2025 IEEE 22nd International Symposium on Biomedical Imaging (ISBI), 1–5. 10.1109/isbi60581.2025.10981062

Le Provost, J.-B., Bartres-Faz, D., Paillere-Martinot, M.-L., Artiges, E., Pappata, S., Recasens, C., Perez-Gomez, M., Bernardo, M., Baeza, I., Bayle, F., & Martinot, J.-L. (2003). Paracingulate sulcus morphology in men with early-onset schizophrenia. The British Journal of Psychiatry: The Journal of Mental Science, 182, 228–232. 10.1192/bjp.182.3.228

Lopez-Persem, A., Verhagen, L., Amiez, C., Petrides, M., & Sallet, J. (2019). The Human Ventromedial Prefrontal Cortex: Sulcal Morphology and Its Influence on Functional Organization. The Journal of Neuroscience: The Official Journal of the Society for Neuroscience, 39(19), 3627–3639. 10.1523/JNEUROSCI.2060-18.2019

Margulies, D. S., Kelly, A. M. C., Uddin, L. Q., Biswal, B. B., Castellanos, F. X., & Milham, M. P. (2007). Mapping the functional connectivity of anterior cingulate cortex. NeuroImage, 37(2), 579–588. 10.1016/j.neuroimage.2007.05.019

Mei, T., Forde, N. J., Floris, D. L., Dell’Acqua, F., Stones, R., Ilioska, I., Durston, S., Moessnang, C., Banaschewski, T., Holt, R. J., Baron-Cohen, S., Rausch, A., Loth, E., Oakley, B., Charman, T., Ecker, C., Murphy, D. G. M., EU-AIMS LEAP group, Beckmann, C. F., … Buitelaar, J. K. (2023). Autism Is Associated With Interindividual Variations of Gray and White Matter Morphology. Biological Psychiatry. Cognitive Neuroscience and Neuroimaging, 8(11), 1084–1093. 10.1016/j.bpsc.2022.08.011

Meredith, S. M., Whyler, N. C. A., Stanfield, A. C., Chakirova, G., Moorhead, T. W. J., Job, D. E., Giles, S., McIntosh, A. M., Johnstone, E. C., & Lawrie, S. M. (2012). Anterior cingulate morphology in people at genetic high-risk of schizophrenia. European Psychiatry: The Journal of the Association of European Psychiatrists, 27(5), 377–385. 10.1016/j.eurpsy.2011.11.004

Nishikuni, K., & Ribas, G. C. (2013). Study of fetal and postnatal morphological development of the brain sulci: Laboratory investigation. Journal of Neurosurgery. Pediatrics, 11(1), 1–11. 10.3171/2012.9.peds12122

Ono, M., Kubik, S., & Abernathey, C. D. (1990). Atlas of the Cerebral Sulci. G. Thieme Verlag. https://play.google.com/store/books/details?id=L7tqAAAAMAAJ

Paus, T., Tomaiuolo, F., Otaky, N., MacDonald, D., Petrides, M., Jason Atlas, Morris, R., & Evans, A. C. (1996). Human Cingulate and Paracingulate Sulci: Pattern, Variability, Asymmetry, and Probabilistic Map. In Cerebral Cortex (Vol. 6, Issue 2, pp. 207–214). 10.1093/cercor/6.2.207

Prigge, M. B. D., Lange, N., Bigler, E. D., King, J. B., Dean, D. C., 3rd, Adluru, N., Alexander, A. L., Lainhart, J. E., & Zielinski, B. A. (2021). A 16-year study of longitudinal volumetric brain development in males with autism. NeuroImage, 236, 118067. 10.1016/j.neuroimage.2021.118067

Ramos Benitez, J., Kannan, S., Hastings, W. L., Parker, B. J., Willbrand, E. H., & Weiner, K. S. (2024). Ventral temporal and posteromedial sulcal morphology in autism spectrum disorder. Neuropsychologia, 195, 108786. 10.1016/j.neuropsychologia.2024.108786

Shim, G., Jung, W. H., Choi, J.-S., Jung, M. H., Jang, J. H., Park, J.-Y., Choi, C.-H., Kang, D.-H., & Kwon, J. S. (2009). Reduced cortical folding of the anterior cingulate cortex in obsessive-compulsive disorder. Journal of Psychiatry & Neuroscience: JPN, 34(6), 443–449. https://www.ncbi.nlm.nih.gov/pubmed/19949720

Team, R. (2020). RStudio: Integrated Development for R. RStudio, PBC, Boston, MA, 2020.

Van Essen, D. C. (2007). 4.16 - Cerebral Cortical Folding Patterns in Primates: Why They Vary and What They Signify. In J. H. Kaas (Ed.), Evolution of Nervous Systems (pp. 267–276). Academic Press. 10.1016/B0-12-370878-8/00344-X

Vogt, B. A., Nimchinsky, E. A., Vogt, L. J., & Hof, P. R. (1995). Human cingulate cortex: surface features, flat maps, and cytoarchitecture. The Journal of Comparative Neurology, 359(3), 490–506. 10.1002/cne.903590310

Watanabe, H., Nakamura, M., Ohno, T., Itahashi, T., Tanaka, E., Ohta, H., Yamada, T., Kanai, C., Iwanami, A., Kato, N., & Hashimoto, R. (2014). Altered orbitofrontal sulcogyral patterns in adult males with high-functioning autism spectrum disorders. Social Cognitive and Affective Neuroscience, 9(4), 520–528. 10.1093/scan/nst016

Welker, W. (1990). Why Does Cerebral Cortex Fissure and Fold? In E. G. Jones & A. Peters (Eds.), Cerebral Cortex: Comparative Structure and Evolution of Cerebral Cortex, Part II (pp. 3–136). Springer US. 10.1007/978-1-4615-3824-0_1

Whittle, S., Chanen, A. M., Fornito, A., McGorry, P. D., Pantelis, C., & Yücel, M. (2009). Anterior cingulate volume in adolescents with first-presentation borderline personality disorder. Psychiatry Research, 172(2), 155–160. 10.1016/j.pscychresns.2008.12.004

Willbrand, E. H., Maboudian, S. A., Elliott, M. V., Kellerman, G. M., Johnson, S. L., & Weiner, K. S. (2024). Variable presence of an evolutionarily new brain structure is related to trait impulsivity. Biological Psychiatry: Cognitive Neuroscience and Neuroimaging. 10.1016/j.bpsc.2024.11.015

Willbrand, E. H., Maboudian, S. A., Kelly, J. P., Parker, B. J., Foster, B. L., & Weiner, K. S. (2023). Sulcal morphology of posteromedial cortex substantially differs between humans and chimpanzees. Communications Biology, 6(1), 1–14. 10.1038/s42003-023-04953-5

Willbrand, E. H., Tsai, Y.-H., Gagnant, T., & Weiner, K. S. (2023). Updating the sulcal landscape of the human lateral parieto-occipital junction provides anatomical, functional, and cognitive insights. In eLife. 10.7554/elife.90451.1

Willbrand, E. H., Voorhies, W. I., Yao, J. K., Weiner, K. S., & Bunge, S. A. (2022). Presence or absence of a prefrontal sulcus is linked to reasoning performance during child development. Brain Structure & Function, 227(7), 2543–2551. 10.1007/s00429-022-02539-1

Yang, D. Y.-J., Beam, D., Pelphrey, K. A., Abdullahi, S., & Jou, R. J. (2016). Cortical morphological markers in children with autism: a structural magnetic resonance imaging study of thickness, area, volume, and gyrification. Molecular Autism, 7, 11. 10.1186/s13229-016-0076-x

Yucel, M., Stuart, G. W., Maruff, P., Velakoulis, D., Crowe, S. F., Savage, G., & Pantelis, C. (2001). Hemispheric and Gender-related Differences in the Gross Morphology of the Anterior Cingulate/Paracingulate Cortex in Normal Volunteers: An MRI Morphometric Study. In Cerebral Cortex (Vol. 11, Issue 1, pp. 17–25). 10.1093/cercor/11.1.17

Yücel, M., Stuart, G. W., Maruff, P., Wood, S. J., Savage, G. R., Smith, D. J., Crowe, S. F., Copolov, D. L., Velakoulis, D., & Pantelis, C. (2002). Paracingulate morphologic differences in males with established schizophrenia: a magnetic resonance imaging morphometric study. Biological Psychiatry, 52(1), 15–23. 10.1016/s0006-3223(02)01312-4

Zhou, Y., Shi, L., Cui, X., Wang, S., & Luo, X. (2016). Functional connectivity of the caudal anterior cingulate cortex is decreased in autism. PloS One, 11(3), e0151879. 10.1371/journal.pone.0151879

Zilles, K., Armstrong, E., Schleicher, A., & Kretschmann, H.-J. (1988). The human pattern of gyrification in the cerebral cortex. In Anatomy and Embryology (Vol. 179, Issue 2, pp. 173–179). 10.1007/bf00304699

