## Supplementary Materials for "Anterior cingulate folding pattern is altered in autism spectrum disorder"


### Authors and affiliations

Ethan H. Willbrand[1,2], Samira A. Maboudian[3,4], Jacob J. Ludwig[1], Kevin S. Weiner[3-5*]

*[1]School of Medicine and Public Health, University of Wisconsin–Madison, Madison, WI USA*
*[2]Medical Scientist Training Program, University of Wisconsin–Madison, Madison, WI USA*
*[3]Helen Wills Neuroscience Institute, University of California, Berkeley, Berkeley, CA*
*[4]Department of Neuroscience, University of California, Berkeley, Berkeley, CA*
*[5]Department of Psychology, University of California, Berkeley, Berkeley, CA*



### Key terms

Anterior cingulate cortex • Autism spectrum disorder • Magnetic resonance imaging • Neuroanatomy • Neurodevelopment • Sulci • Cortical folding

**Supplementary Methods**

**Participants**

We leveraged data from the Autism Brain Imaging Data Exchange (ABIDE; (Di Martino et al., 2014), which were used in our previous work in other cortical regions (Ramos Benitez et al., 2024). Given limitations of this dataset resulting from females being historically underdiagnosed for ASD and less present in the ABIDE sample, as well as the large developmental age range of the ABIDE samples (childhood through young adulthood), we specifically selected participants that were male and less than 20 years old. In total, we randomly selected 100 participants fitting these criteria (50 participants in each group). We also ensured that the NT and ASD groups were age matched (NT: 10.02 ± 2.03 years old; ASD: 9.32 ± 2.73 years old; $t(98) = -1.45$, $p = 0.15$).

**Neuroanatomical data**

***Imaging Acquisition:*** All participants who participated in the study completed a mock scanning session in order to become acclimated to the scanning environment. Standard T1-weighted MPRAGE anatomical scans (TR = 2530 ms, TE = 3.25 ms, 1.3 × 1 × 1.3 $mm^3$ voxels) were acquired for cortical morphometric analyses in these participants using a 3T Siemens Allegra MRI scanner at New York University Langone Medical Center, the Center for Functional and Molecular Imaging at Georgetown University, and Stanford School of Medicine Richard M. Lucas Service Center for Imaging.

***Cortical Surface Reconstruction:*** Cortical surface reconstructions were generated for each participant from their T1-weighted image using the standard FreeSurfer pipeline (v6.0; (Dale et al., 1999; Fischl et al., 1999). The manual identification of the PCGS was performed on each cortical surface in native space.

***Paracingulate Sulcus Identification, Classification Criteria, and Morphological Feature Extraction:*** To analyze the organization of the ACC sulci, this study examined the presence or absence of the PCGS, a tertiary sulcus located dorsally to and parallel with the CGS (Paus et al., 1996; Yucel et al., 2001). As is standard in studies examining PCGS presence (Amiez et al., 2018; Garrison et al., 2015; Harper et al., 2022; Ono et al., 1990; Willbrand et al., 2024), a binary classification was employed to assess PCGS presence in each hemisphere, with the sulcus considered "present" if it measured at least 20 mm in length and "absent" if no horizontal sulcal element extended parallel to the CGS for at least this length. The PCGS also had to have a depth of at least 4 mm. Sub-classification of PCGS presence (i.e., distinguishing between “prominent” and “present”) was not analyzed as it is not a consistent measure and varies with demographic features (Cachia et al., 2021; Clark et al., 2010; Harper et al., 2022, 2023; Leonard et al., 2009). We also classified asymmetries in PCGS presence using standard terminology (Paus et al., 1996; Yucel et al., 2001). An asymmetrical PCGS was defined by a difference in presence between hemispheres (e.g., present in the left, absent in the right). Symmetrical patterning required the PCGS to be similarly classified across hemispheres (both present or both absent). The PCGS was assessed by trained raters (EHW, JJL, SAM). 30 hemispheres were independently identified by the three raters to calculate inter-rater agreement, which was near perfect (Cohen’s $\kappa = 0.93$). See **Figure 1** for examples of PCGS presence/absence and asymmetry.

To acquire quantitative morphological features of the PCGS, the PCGS was manually defined in each hemisphere as a .label file using FreeSurfer’s tksurfer tools. The length (in mm) was calculated as the longest geodesic distance between any boundary vertices involving the .label file using algorithms implemented in the pycortex python package (Gao et al., 2015). Disconnected sulcal pieces were combined to determine total length excluding gyral components. PCGS depth was calculated from the sulcal fundus to the outer pial surface using a modified algorithm building on the FreeSurfer pipeline (Madan, 2019). We also extracted cortical thickness (CT) mean and standard deviation (STD) values using the built-in mris_anatomical_stats FreeSurfer function (Fischl & Dale, 2000), given prior literature showing alterations in CT in the vicinity of the PCGS at the group level (Abell et al., 1999) and CT STD in tertiary sulci in ASD (Ammons et al., 2021; Ramos Benitez et al., 2024).

***Calculating the amount of cortex buried in ACC:*** To investigate whether differences in folding observed at the individual sulcal level were present at the regional level, we measured the amount of ACC buried within sulci. To test this, we identified relevant anatomical regions comprising ACC using the Destrieux parcellation (Destrieux et al., 2010). Specifically, ACC was defined by combining four Destrieux regions (anterior, middle-anterior, and middle-posterior parts of the cingulate gyrus and sulcus, and superior frontal gyrus). These regions were extracted from the Destrieux annotation, converted into individual labels, and merged into a single "ACC ROI" label file using FreeSurfer's mri_annot2label and mri_mergelabels functions.

To quantify the portions of cortex composed of sulci, we used FreeSurfer's .curv file, which distinguishes sulcal from gyral regions on cortical surfaces (Dale et al., 1999). A "sulci ROI" label file was generated by thresholding the .curv file to include all vertices with values greater than zero (i.e., the sulcal regions), using the mri_binarize function in FreeSurfer. Finally, to assess the proportion of ACC composed of sulcal cortex, we computed the Dice coefficient which quantifies the overlap between the ACC ROI and the sulci ROI as utilized in prior work (Miller et al., 2021; Ramos Benitez et al., 2024; Willbrand, Bunge, et al., 2023; Willbrand et al., 2022; Willbrand, Maboudian, et al., 2023):

$$DICE\,(X,Y) \;=\; \frac{2|X \cap Y|}{|X| + |Y|}$$

### Quantification and Statistical Analysis

All statistical tests were implemented in R (v4.1.2). First, for each group separately, chi-squared ($\chi 2$) tests were used to test for intra- and inter-hemispheric differences in PCGS incidence, and

McNemar's test for symmetry was used to test for PCGS asymmetry. Second, to test whether the PCGS incidence/asymmetry related to diagnosis, we ran binomial logistic regression general linear models (GLMs) for PCGS patterning [0 (present or symmetric), 1 (absent or asymmetric)] with diagnostic group (ASD, NT) as the primary factor. GLM effect size was quantified as the Odds ratio (OR). We internally validated any significant diagnosis-related effect by bootstrapping the OR with 1,000 resamples. To assess the relationship between diagnosis and quantitative PCGS morphology, we ran linear mixed-effects models (LMEs) with predictors of diagnostic group (ASD, NT) and hemisphere (left, right) for the four quantitative morphological features of interest (PCGS length, PCGS depth, and PCGS CT mean and STD). Finally, to determine whether the amount of ACC buried in sulci differed by diagnosis and PCGS presence, we implemented an LME with predictors of diagnostic group (ASD, NT), hemisphere (left, right), and PCGS presence (present, absent). In all analyses, we controlled for age and scanner site. In the quantitative morphological analyses, the models all controlled for the relevant hemisphere-wide variable (hemispheric surface area, max depth, and CT mean and STD, respectively).

**Supplementary Results**

**Paracingulate sulcal morphology does not differ in autism spectrum disorder**

We then examined whether quantitative morphological features of the PCGS (length, depth, and CT mean and STD) differed between the two groups by implementing LMEs with predictors of group and hemisphere (controlling for age, scanner site, and the respective hemispheric-level morphological variable). None of these analyses revealed any significant main effects of group or interactions (all *p*s > 0.05; **Supplementary Figure 1**). We did observe a significant effect of hemisphere on PCGS mean cortical thickness ($F(1, 51) = 10.32$, $p = 0.002$), such that the gray matter in left PCGS were thicker on average than right PCGS across groups (**Supplementary Figure 1D**). There were no other hemispheric differences observed for the other three quantitative morphological features (all *p*s > 0.05; **Supplementary Figure 1**).

**The amount of anterior cingulate cortex buried in sulci does not differ in autism spectrum disorder**

Finally, we compared how much of the ACC was buried within sulci (via the Dice coefficient overlap between a parcellation of ACC and the amount of cortex identified as sulci; **Materials and Methods**; **Supplementary Figure 2A**) between groups and whether this was impacted by the presence or absence of the PCGS. Consistent with prior work using other methods and in other regions (Ramos Benitez et al., 2024; Van Essen, 2007; Vogt et al., 1995; Willbrand, Maboudian, et al., 2023; Zilles et al., 1988), the majority of human ACC is buried within sulci (overall mean ± sd: 61.25 ± 0.01%; **Supplementary Figure 2B**). Next, a LME with predictors of group (ASD, NT), hemisphere (left, right), and PCGS presence (present, absent; controlling for age and scanner site) revealed two findings. First, there were no main effect or interactions with group (all *p*s > 0.05; **Supplementary Figure 2B**). Second, there was a main effect of PCGS presence ($F(1, 94) = 4.55$, $p = 0.035$), such that an ACC containing a PCGS had more surface buried within sulci (**Supplementary Figure 2C**).

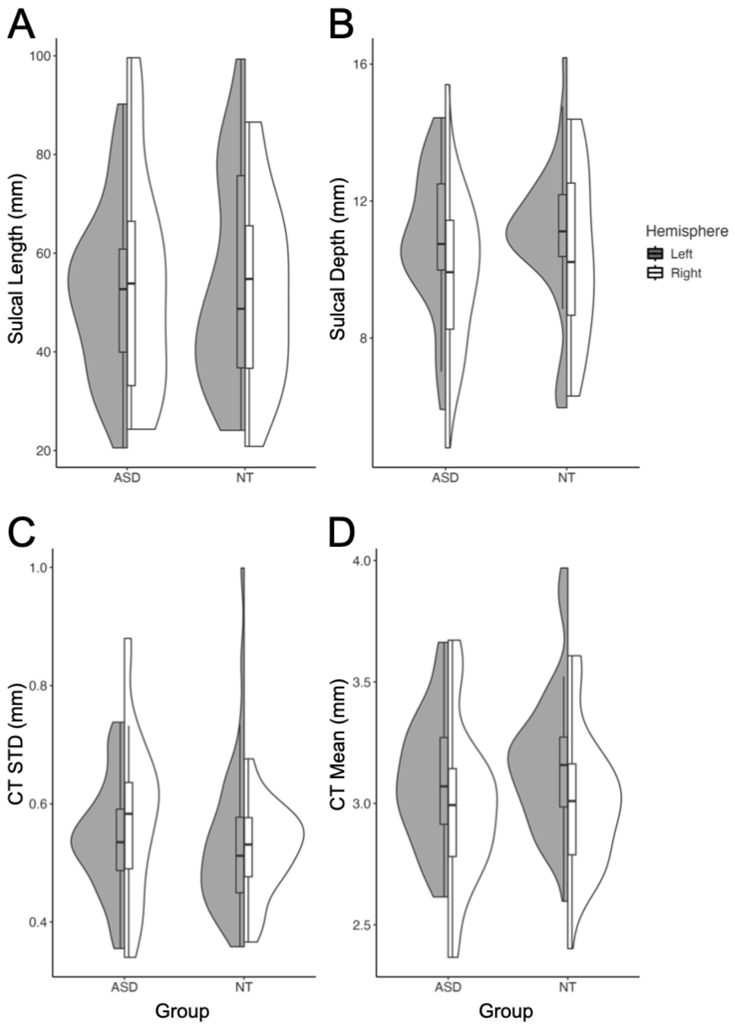


**Supplementary Figure 1. Quantitative morphological features of the PCGS do not differ between ASD and NT individuals. (A)** Split violin plots (box plot and kernel density estimate) visualizing PCGS sulcal length (in mm) as a function of diagnosis (x-axis) and hemisphere (colors; see key). **(B)** Same format as (A), but for PCGS depth (in mm). **(C)** Same format as (A), but for PCGS CT STD (in mm). **(D)** Same format as (A), but for PCGS CT mean (in mm). Abbreviations are as follows: autism spectrum disorder (ASD), cortical thickness (CT), diagnosis (DX), neurotypical (NT), paracingulate sulcus (PCGS), standard deviation (STD).

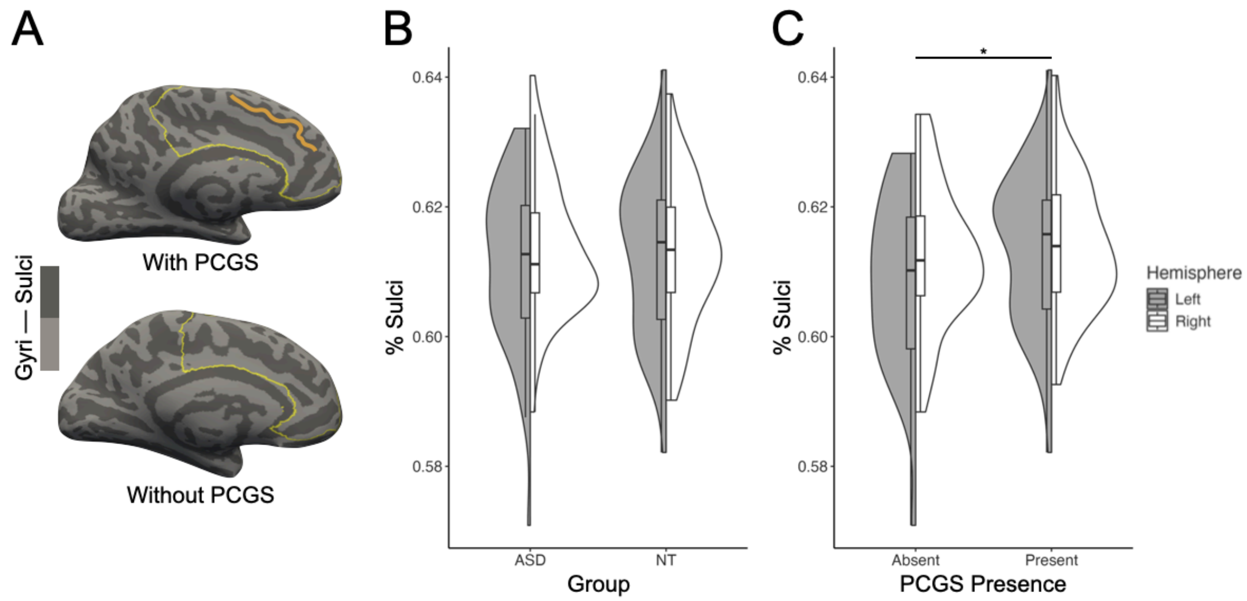


**Supplementary Figure 2. The amount of ACC buried in sulci does not differ between ASD and NT individuals. (A)** Example inflated FreeSurfer cortical surface reconstructions with (top) and without (bottom) a PCGS. The PCGS is highlighted in orange as in Fig. 1. Sulci are dark gray and gyri are light gray. The area defined as ACC (via the Destrieux parcellation) (Destrieux et al., 2010) is outlined in yellow. The PCGS is colored according to Figure 1. **(B)** Split violin plots (box plot and kernel density estimate) visualizing the percent of ACC buried in sulci (Dice coefficient; values are out of 1) as a function of diagnosis (x-axis) and hemisphere (colors; see key). **(C)** Same format as (B), but for PCGS presence (x-axis). Abbreviations are as follows: autism spectrum disorder (ASD), diagnosis (DX), neurotypical (NT), paracingulate sulcus (PCGS). Asterisks indicate the following *p*-value thresholds: * $p < 0.05$.

**Supplementary References**


Abell, F., Krams, M., Ashburner, J., Passingham, R., Friston, K. J., Frackowiak, R. S. J., Happé, F., Frith, C., & Frith, U. (1999). The neuroanatomy of autism: a voxel-based whole brain analysis of structural scans. *Neuroreport*, *10*(8), 1647–1651. https://doi.org/10.1097/00001756-199906030-00005

Amiez, C., Wilson, C. R. E., & Procyk, E. (2018). Variations of cingulate sulcal organization and link with cognitive performance. *Scientific Reports*, *8*(1), 1–13. https://doi.org/10.1038/s41598-018-32088-9

Ammons, C. J., Winslett, M.-E., Bice, J., Patel, P., May, K. E., & Kana, R. K. (2021). The mid-fusiform sulcus in autism spectrum disorder: Establishing a novel anatomical landmark related to face processing. *Autism Research: Official Journal of the International Society for Autism Research*, *14*(1), 53–64. https://doi.org/10.1002/aur.2425

Cachia, A., Borst, G., Jardri, R., Raznahan, A., Murray, G. K., Mangin, J.-F., & Plaze, M. (2021). Towards Deciphering the Fetal Foundation of Normal Cognition and Cognitive Symptoms From Sulcation of the Cortex. *Frontiers in Neuroanatomy*, *15*, 712862. https://doi.org/10.3389/fnana.2021.712862

Clark, G. M., Mackay, C. E., Davidson, M. E., Iversen, S. D., Collinson, S. L., James, A. C., Roberts, N., & Crow, T. J. (2010). Paracingulate sulcus asymmetry; sex difference, correlation with semantic fluency and change over time in adolescent onset psychosis. *Psychiatry Research*, *184*(1), 10–15. https://doi.org/10.1016/j.pscychresns.2010.06.012

Dale, A. M., Fischl, B., & Sereno, M. I. (1999). Cortical surface-based analysis. I. Segmentation and surface reconstruction. *NeuroImage*, *9*(2), 179–194. https://doi.org/10.1006/nimg.1998.0395

Destrieux, C., Fischl, B., Dale, A., & Halgren, E. (2010). Automatic parcellation of human cortical gyri and sulci using standard anatomical nomenclature. *NeuroImage*, *53*(1), 1–15. https://doi.org/10.1016/j.neuroimage.2010.06.010

Di Martino, A., Yan, C.-G., Li, Q., Denio, E., Castellanos, F. X., Alaerts, K., Anderson, J. S., Assaf, M., Bookheimer, S. Y., Dapretto, M., Deen, B., Delmonte, S., Dinstein, I., Ertl-Wagner, B., Fair, D. A., Gallagher, L., Kennedy, D. P., Keown, C. L., Keysers, C., … Milham, M. P. (2014). The autism brain imaging data exchange: towards a large-scale evaluation of the intrinsic brain architecture in autism. *Molecular Psychiatry*, *19*(6), 659–667. https://doi.org/10.1038/mp.2013.78

Fischl, B., & Dale, A. M. (2000). Measuring the thickness of the human cerebral cortex from magnetic resonance images. *Proceedings of the National Academy of Sciences of the United States of America*, *97*(20), 11050–11055. https://doi.org/10.1073/pnas.200033797

Fischl, B., Sereno, M. I., & Dale, A. M. (1999). Cortical surface-based analysis. II: Inflation, flattening, and a surface-based coordinate system. *NeuroImage*, *9*(2), 195–207. https://doi.org/10.1006/nimg.1998.0396

Gao, J. S., Huth, A. G., Lescroart, M. D., & Gallant, J. L. (2015). Pycortex: an interactive surface visualizer for fMRI. *Frontiers in Neuroinformatics*, *9*, 23. https://doi.org/10.3389/fninf.2015.00023

Garrison, J. R., Fernyhough, C., McCarthy-Jones, S., Haggard, M., Australian Schizophrenia Research Bank, & Simons, J. S. (2015). Paracingulate sulcus morphology is associated with hallucinations in the human brain. *Nature Communications*, *6*, 8956. https://doi.org/10.1038/ncomms9956

Harper, L., de Boer, S., Lindberg, O., Lätt, J., Cullen, N., Clark, L., Irwin, D., Massimo, L., Grossman, M., Hansson, O., Pijnenburg, Y., McMillan, C. T., & Santillo, A. F. (2023). Anterior cingulate sulcation is associated with onset and survival in frontotemporal dementia. *Brain Communications*, *5*(5), fcad264. https://doi.org/10.1093/braincomms/fcad264

Harper, L., Lindberg, O., Bocchetta, M., Todd, E. G., Strandberg, O., van Westen, D., Stomrud, E., Landqvist Waldö, M., Wahlund, L.-O., Hansson, O., Rohrer, J. D., & Santillo, A. (2022).

Prenatal Gyrification Pattern Affects Age at Onset in Frontotemporal Dementia. *Cerebral Cortex* . https://doi.org/10.1093/cercor/bhab457

Leonard, C. M., Towler, S., Welcome, S., & Chiarello, C. (2009). Paracingulate asymmetry in anterior and midcingulate cortex: sex differences and the effect of measurement technique. *Brain Structure & Function*, *213*(6), 553–569. https://doi.org/10.1007/s00429-009-0210-z

Madan, C. R. (2019). Robust estimation of sulcal morphology. *Brain Informatics*, *6*(1), 5. https://doi.org/10.1186/s40708-019-0098-1

Miller, J. A., Voorhies, W. I., Lurie, D. J., D'Esposito, M., & Weiner, K. S. (2021). Overlooked Tertiary Sulci Serve as a Meso-Scale Link between Microstructural and Functional Properties of Human Lateral Prefrontal Cortex. *The Journal of Neuroscience: The Official Journal of the Society for Neuroscience*, *41*(10), 2229–2244. https://doi.org/10.1523/JNEUROSCI.2362-20.2021

Ono, M., Kubik, S., & Abernathey, C. D. (1990). *Atlas of the Cerebral Sulci*. G. Thieme Verlag. https://play.google.com/store/books/details?id=L7tqAAAAMAAJ

Paus, T., Tomaiuolo, F., Otaky, N., MacDonald, D., Petrides, M., Jason Atlas, Morris, R., & Evans, A. C. (1996). Human Cingulate and Paracingulate Sulci: Pattern, Variability, Asymmetry, and Probabilistic Map. In *Cerebral Cortex* (Vol. 6, Issue 2, pp. 207–214). https://doi.org/10.1093/cercor/6.2.207

Ramos Benitez, J., Kannan, S., Hastings, W. L., Parker, B. J., Willbrand, E. H., & Weiner, K. S. (2024). Ventral temporal and posteromedial sulcal morphology in autism spectrum disorder. *Neuropsychologia*, *195*, 108786. https://doi.org/10.1016/j.neuropsychologia.2024.108786

Van Essen, D. C. (2007). 4.16 - Cerebral Cortical Folding Patterns in Primates: Why They Vary and What They Signify. In J. H. Kaas (Ed.), *Evolution of Nervous Systems* (pp. 267–276). Academic Press. https://doi.org/10.1016/B0-12-370878-8/00344-X

Vogt, B. A., Nimchinsky, E. A., Vogt, L. J., & Hof, P. R. (1995). Human cingulate cortex: surface features, flat maps, and cytoarchitecture. *The Journal of Comparative Neurology*, *359*(3),

490–506. https://doi.org/10.1002/cne.903590310

Willbrand, E. H., Bunge, S. A., & Weiner, K. S. (2023). Neuroanatomical and Functional Dissociations between Variably Present Anterior Lateral Prefrontal Sulci. *Journal of Cognitive Neuroscience*, *35*(11), 1846–1867. https://doi.org/10.1162/jocn_a_02049

Willbrand, E. H., Maboudian, S. A., Elliott, M. V., Kellerman, G. M., Johnson, S. L., & Weiner, K. S. (2024). Variable presence of an evolutionarily new brain structure is related to trait impulsivity. *Biological Psychiatry: Cognitive Neuroscience and Neuroimaging*. https://doi.org/10.1016/j.bpsc.2024.11.015

Willbrand, E. H., Maboudian, S. A., Kelly, J. P., Parker, B. J., Foster, B. L., & Weiner, K. S. (2023). Sulcal morphology of posteromedial cortex substantially differs between humans and chimpanzees. *Communications Biology*, *6*(1), 1–14. https://doi.org/10.1038/s42003-023-04953-5

Willbrand, E. H., Parker, B. J., Voorhies, W. I., Miller, J. A., Lyu, I., Hallock, T., Aponik-Gremillion, L., Koslov, S. R., Null, N., Bunge, S. A., Foster, B. L., & Weiner, K. S. (2022). Uncovering a tripartite landmark in posterior cingulate cortex. *Science Advances*, *8*(36), eabn9516. https://doi.org/10.1126/sciadv.abn9516

Yucel, M., Stuart, G. W., Maruff, P., Velakoulis, D., Crowe, S. F., Savage, G., & Pantelis, C. (2001). Hemispheric and Gender-related Differences in the Gross Morphology of the Anterior Cingulate/Paracingulate Cortex in Normal Volunteers: An MRI Morphometric Study. In *Cerebral Cortex* (Vol. 11, Issue 1, pp. 17–25). https://doi.org/10.1093/cercor/11.1.17

Zilles, K., Armstrong, E., Schleicher, A., & Kretschmann, H.-J. (1988). The human pattern of gyrification in the cerebral cortex. In *Anatomy and Embryology* (Vol. 179, Issue 2, pp. 173–179). https://doi.org/10.1007/bf00304699
